# Drivers of change in the realised climatic niche of terrestrial mammal species

**DOI:** 10.1101/2020.03.12.985374

**Authors:** Di Marco Moreno, Michela Pacifici, Luigi Maiorano, Carlo Rondinini

## Abstract

The breadth of a species’ climatic niche is an important ecological trait that allows adaptation to climate change, but human activities drive niche erosion. Life-history traits, such as dispersal ability and reproductive speed, instead allow species to cope with climate change. But how do these characteristics act in combination with human pressure to determine niche change? Here we investigate the patterns and drivers of change in the realised climatic niche of 589 terrestrial mammal species. Our goal is to disentangle the impacts of humans, climate change, and life history. We calibrated the past and present climatic niches of each species by considering past climatic conditions (Mid Holocene) within their pre-human impact distributions, and current climatic conditions within the current distributions. Depending on the relationship between past and current niche, we defined four categories of change: “shrink”, “shift”, “stable”, and “expand”. We found over half of the species in our sample have undergone niche shrink, while only 15-18% of species retained a stable niche. After controlling for biogeography, climatic factors were the strongest correlates of species niche change, followed by anthropogenic pressure and species’ life history. Factors that increased the probability of niche shrink include: overall climatic instability in the area (both intermediate or high), large body mass, long gestation time, highly carnivorous or herbivorous diets, historical land-use change, and current human population density. We identified the conditions under which species are less likely to maintain their niche breadth, potentially losing adaptation capacity under climate change. Species with these characteristics require interventions that facilitate natural dispersal or assisted colonisation, to survive to rapidly changing climates.

## Introduction

The breadth of a species’ niche - the set of environmental conditions in which the species can persist (Peterson et al. 2011) - is an important ecological trait that allows adaptation to environmental change (Thuiller et al. 2005; Pacifici et al. 2015). Niche breadth is a key correlate of species sensitivity to future climate change (Swihart et al. 2003; Thuiller et al. 2005; Chown et al. 2010), and is usually assessed by relating the observed occurrences of species to their respective climate. This implies looking at species’ realised niches, rather than their fundamental ones (Peterson et al. 2011). In fact, analysing realised niches is a well-established technique to identify differences in species’ ecology (Olalla-Tárraga et al. 2011; Mahon et al. 2016), predict the potential spread of invasive species (Liu et al. 2017), and project past and future changes in species distributions (Maiorano et al. 2013; Visconti et al. 2016).

While the roles of human threats as drivers of species decline and extinction have been often demonstrated (Johnson et al. 2017; Pacifici et al. 2017; Di Marco et al. 2018), their role as drivers of niche erosion has proven more difficult to quantify (Pearman et al. 2008). Yet this is a critical element to consider, because disregarding the effect of human modifications of species realized niches might result in biased estimates of the future impact of climate change (Faurby & Araújo 2018). Some evidence of how humans have altered species niche is already available, despite uncertainty in past information on species distribution, climatic conditions, and human pressure (Walther et al. 2005). For example, analyses on the distribution range of the giraffe (*Giraffa Camelopardalis*) and African elephant (*Loxodonta Africana*) in the last 150 years show a reduction in their climatic niche as a consequence of poaching, fragmentation, and conflicts (Martínez-Freiría et al. 2015).

Threats such as overexploitation, habitat loss and fragmentation, or invasive species have been recognised as global drivers of species decline in recent centuries (Hoffmann et al. 2010; Maxwell et al. 2016). More recently, substantial attention has been devoted to the emerging threat of climate change, with effects that might become soon predominant over those of already established threats (Newbold 2018; Di Marco et al. 2019). Climate change is recognised to have potential magnifying effects on biodiversity decline in the absence of adaptation and coping mechanisms (Bellard et al. 2012; Mantyka-Pringle et al. 2015; Visconti et al. 2016). Yet species might be able to tolerate changing climates to some extent, depending on their characteristics (Adrian et al. 2006; Jiguet et al. 2007; Urban et al. 2014; Santini et al. 2016; Pacifici et al. 2017). Life-history traits, such as dispersal ability and reproductive speed for example, have been hypothesized to play a central role in determining the sensitivity of species to climate change and their ability to cope with it (Dawson 2011). Evolutionary adaptation might also allow species to cope with changing climate (Hoffmann & Sgró 2011), even if it is unclear whether this mechanism is compatible with the pace of current climate change (Loarie et al. 2009). But how do these mechanisms act in combination with human pressure to determine change in species climatic niches?

Here we investigate the patterns and drivers of change in the realised climatic niche of terrestrial mammals. Our goal is to disentangle the impacts of humans, climate change, and life history on species climatic niches. Separating intrinsic and extrinsic vulnerability of species to niche change, as well as the role of direct and indirect human pressure, is essential to understand which species are unlikely to adapt to future climatic conditions. We focus our analysis on terrestrial mammals, a data-rich group compared to other taxa, given the availability of distribution data for all species, both at present (IUCN 2018) and before human impact took place (Faurby & Svenning 2015). Terrestrial mammals make fundamental contributions to key ecological processes such as predation, herbivory, and seed dispersal, but are facing high risk of extinction (Fragoso et al. 2003; Soulé & Estes 2003; Pringle et al. 2007; Hoffmann et al. 2011). Their ability to adapt to rapidly changing climate (or lack thereof) is an essential element to consider when forecasting future extinction rates and defining appropriate conservation measures (Pacifici et al. 2017).

## Methods

### Species data

We focused our analyses on 589 terrestrial mammal species (Table S1), representing all species which are known to have changed their geographic distribution in response to human pressure, and have been assessed in the Red List of the International Union for Conservation of Nature (IUCN). Selecting these species allowed us to disentangle the relative impact of climate change (within species’ natural ranges) from that of direct human influence on species’ distributions (Faurby & Svenning 2015). We used species distributions referring to the present day, and those assumed to represent species’ natural ranges (i.e. before human impact modified them). We retrieved present distributions from the IUCN Red List (IUCN 2018) and pre-impact distributions from the PHYLACINE dataset (Faurby & Svenning 2015; Faurby et al. 2018). All ranges were considered at a spatial resolution of 1 degree (roughly 110 km x 110 km at the equator), which is the native resolution in the PHYLACINE database.

We collected life-history and ecological traits of species that are potentially correlated to change in their realised climatic niches. We considered the following variables: species biogeographic domains (Olson et al. 2001), percentage of vertebrate/invertebrate/plat diet (Faurby et al. 2018), body mass (Faurby et al. 2018), gestation length (Jones et al. 2009; Tacutu et al. 2013), and interbirth interval (Jones et al. 2009; Tacutu et al. 2013). Missing data for gestation length and interbirth interval were imputed from other life-history traits and phylogeny, using the R package “missForest” (Stekhoven & Bühlmann 2012) and following the procedure of Penone et al. (2014). During imputation process, we represented species phylogeny by extracting phylogenetic eigenvectors (Diniz-Filho et al. 1998) from the PHYLACINE dataset (Faurby et al. 2018). That phylogeny was derived using a hierarchical Bayesian approach with a posterior distribution of 1,000 trees, which represent uncertainties in topology and branch lengths. We extracted 10 random trees from the phylogeny and re-ran our data imputation process using each of the trees, to test the sensitivity of our imputation to phylogenetic uncertainty. We also verified whether directly including phylogenetic relationships improved the performance of our niche models, using phylogenetic eigenvectors as model predictors (see below).

We also included anthropogenic drivers of change in species niches. We quantified the levels of human pressure to which species were exposed through time, by accounting for past and current levels of human encroachment within species’ natural ranges (pre-impact distributions). We measured both human population density and the amount of agricultural land within each species’ range. We derived population densities and land-use data for the years 4,000 BC and 2017 AD, from the HYDE dataset at their original 5 arc-minutes resolution (Goldewijk et al. 2017). This corresponds to the Mid Holocene climatic period from Worldclim (Hijmans et al. 2005), which we used for representing past species niches (see below). We measured different percentiles of the distribution of these pressures within species ranges, and selected the percentile leading to highest predictive importance of the variables (Di Marco et al. 2015): 95% for past land use, 50% for current land use, 75% for past human density, and 25% for current human density.

### Representing species climatic niches

We calibrated the past and present climatic niches of each species by considering the past climatic conditions within their pre-impact range and the current conditions in the current range. We considered past climatic conditions in the Mid Holocene (MID; ca. year 4,000 BC) as obtained by the IPSL-CM5A-LR and the MPI-ESM-P general circulation models (GCMs). We averaged the results based on those two GCMs to account for uncertainty in past climatic projections. We also considered conditions at an earlier period, the Last Inter-Glacial (LIG; ca. 130,000 years ago) as a sensitivity test. A set of 10 bioclimatic variables were extracted from the Worldclim dataset (Hijmans et al. 2005), previously identified for their ability to model mammal species’ climatic preferences (Visconti et al. 2016): Annual Mean Temperature, Mean Temperature of Wettest Quarter, Mean Temperature of Driest Quarter, Mean Temperature of Warmest Quarter, Mean Temperature of Coldest Quarter, Annual Precipitation, Precipitation of Wettest Quarter, Precipitation of Driest Quarter, Precipitation of Warmest Quarter, Precipitation of Coldest Quarter. We extracted climatic conditions within the pre-impact and current distribution range of each species at a resolution of 30 arc-seconds (approximately 1 km at the equator), which is the common resolution among the various climatic datasets we analysed. We represented fine-scale climatic conditions throughout each species’ range, by sampling the centroid of each 2.5 arc-minutes grid cell (approximately 5 km x 5 km at the equator) within the coarse species ranges, as a compromise between spatial coverage and computational feasibility. We extracted the climatic characteristics for all sampled pixels and analysed them using a principal component analysis (PCA) approach. This way we represented the combination of relatively fine-scaled climatic conditions that a species experienced within its broad distribution range through time.

The delineation of species niches was done using the software R (R Core Team 2018) and the package “ecospat” (Di Cola et al. 2017). We treated the pre-impact and current species distributions in a similar way to how native and non-native distributions are treated when investigating niche change for invasive species. We followed Broennimann et al. (2012) in defining a gridded ecological niche space for each species, delimited by the two major axes of a PCA built on the above-listed bioclimatic variables. We defined such environmental space by using past and present climate within each species’ biogeographic domain as the reference climatic regions, and the climate registered within pre-impact and current species distribution as proxy of species realised niches. This implies each species is associated to a “study region” that represents its biogeographical domain. We projected the PCA scores of the past and current climate experienced by the species onto the gridded ecological space, to define smoothed density of occurrences using a kernel density function.

We represented the past and current niches as the polygons encompassing 95% of the gridded occurrences, respectively around the pre-impact occurrences and the current occurrences. We classified categories of change in the realised climatic niches of terrestrial mammal species by considering the relative size and position of the niche polygons in the gridded environmental space (Fig. 1). In particular, we defined four categories of niche change: “shrink”, when a species’ niche has reduced over time; “shift”, when a species niche has changed position without substantial reduction in its variability; “stable”, when a species niche has not substantially reduced or shifted; “expand”, when a species’ niche has increased in size over time.

**Figure 1.**
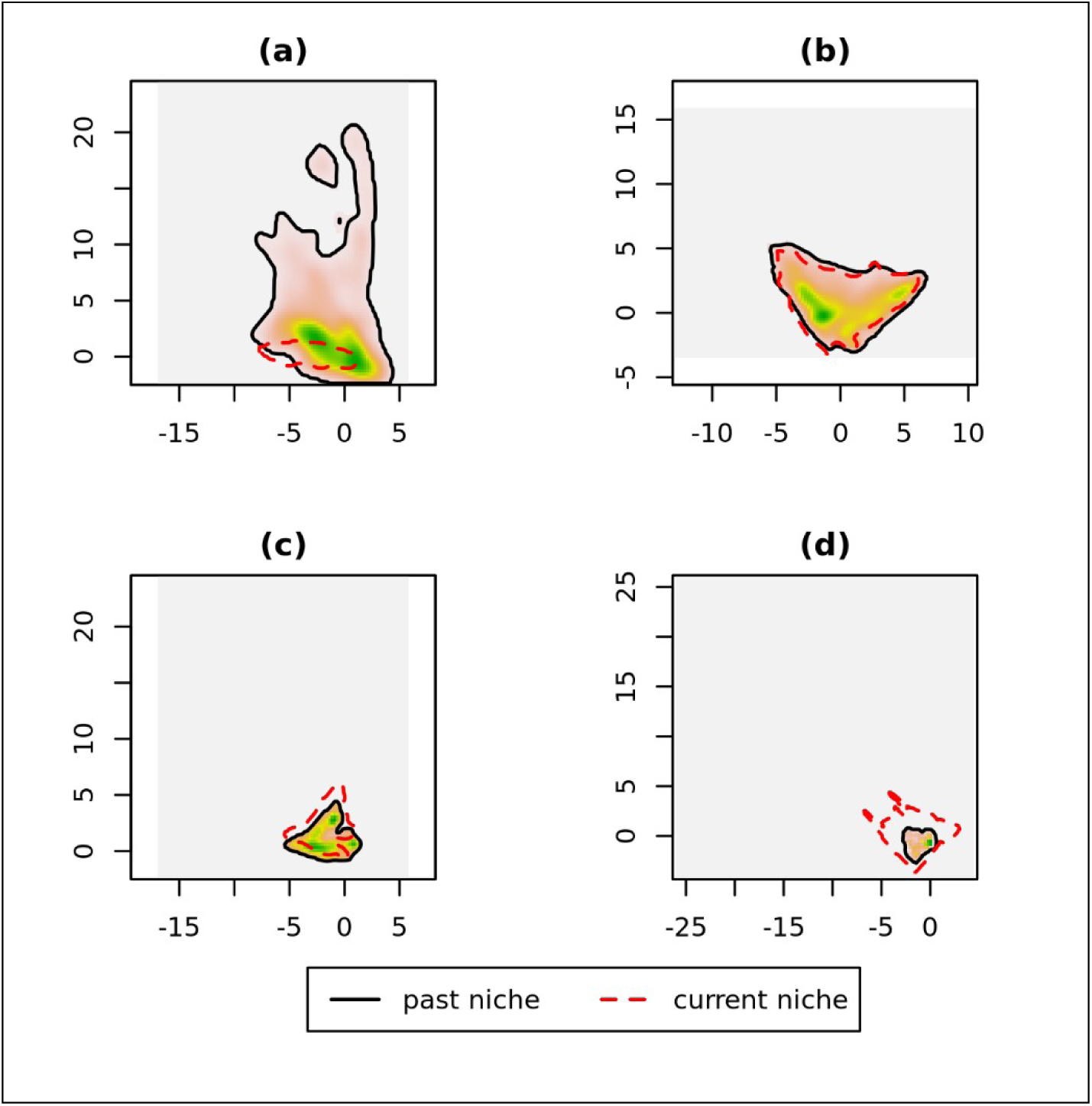
Categories of change in the realised climatic niches of terrestrial mammal species, derived by comparison of past climate in pre-human impact distribution (black solid line) and current climate in current distribution (red dashed line). The density of species distribution within the past niche is represented as an orange-to-green gradient. The four panels represent: a) an example of niche “shrink”, the Ethiopian wolf (Canis simensis); b) an example of niche “stability”, the Spectacled Bear (Tremarctos ornatus); c) an example of niche “shift”, the Eastern Red Forest Rat (Nesomys rufus); d) an example of niche “expansion”, the Coyote (Canis latrans).

We defined thresholds of tolerance below which niche changes were considered minimal, and the species classified as “stable”; this way we prevented the model from being over-sensitive to data uncertainty (e.g., in terms of past climate and species distributions). We tested tolerance thresholds of 5%, 10%, and 20% to separate niche stability from niche change, both in terms of position shift and in terms of shrink/expansion, and to separate niche shift from niche expansions and shrinks.

### Modelling change in species climatic niche

We run multinomial logistic regression models to predict the probability of species to be assigned to a given class of niche change, using the R package ‘nnet’ (Venables & Ripley 2002). The same model structure was repeated under different past climate scenarios (in terms of GCM and time period). We used the class “shrink” as the reference level in the models. We included all the above-described intrinsic and extrinsic variables as model predictors, after verifying that these are not collinear with each other (Pearson’s r < 0.7). All continuous variables were scaled to improve comparability of model’s coefficients. In order to disentangle the effect of regional climate change from that of other drivers influencing the dynamics of a species’ niche, we also measured the overall climatic stability within species pre-impact ranges. We did so by measuring the proportion of past climatic space that is retained in the present, within the same PCA gridded ecological space used to define species niches. We used this metric of overall climate stability as one of the predictors in our model.

We measured the model’s fit using Nagelkerke pseudo-R^2^, and evaluated the model’s performance using a leave-one-out validation approach. The validation routine was performed by iteratively excluding one species at a time, and then using the model calibrated on all other species to predict the probability that the left-out species belongs to each of the four classes of niche change. We compared the predicted class probabilities with the original (observed) class of each species. We measured the model’s classification accuracy with three different metrics. First, we defined a “predicted class” for each species, as the class with the highest assigned probability by the model. Second, for each species we ranked the predicted niche classes from the most probable to the least probable, and measured the rank of the observed class. Third, for each species we measured the difference between the predicted probability of the observed category and that of the most probable category according to the model (δ-prediction). This value would be zero if the observed class is also the one with the highest predicted probability, and >0 otherwise; as an example, a species with a “stable” niche for which the models assigns a 50% chance of niche “shift” and a 40% chance of niche “stable” has a δ-prediction of 10%. We estimated both the overall classification accuracy, across all species, and the accuracy for species in each separate category of niche change.

We estimated the models’ coefficients and their statistical significance, and we represented the relationship between key predictors in our model and the probability of being in a given class of niche change. To represent the latter relationships, we produced partial effect plots that represent the effect of one variable (e.g. body mass) on the response (e.g probability of the species to belong to the category “stable niche”) while holding all other variables constant. Finally, we compared our results from the multinomial model to those obtained with a Random Forest model (a non-parametric machine-learning technique), using the R package ‘randomForest’ (Liaw & Wiener 2002).

## Results

### Model’s ability to classify change in species niche

The combination of MID climate (under the IPSL-CM5A-LR General Circulation Model) and 20% tolerance threshold, resulted in the best model’s performances (Table S2). Under those settings, the model had good overall classification accuracy (59% species correctly assigned to their observed niche classes) and a lower class-averaged accuracy (43%), which was still the highest value across all model settings. In fact, under any climatic scenario, using higher tolerance thresholds led to an increase in the variance explained by the model, a decrease in the overall prediction accuracy (across all species), and an increase in the class-averaged prediction accuracy. The mean prediction rank of the observed class was 1.64, while the average δ-prediction value was 15% (Table S3). The most numerous class, niche shrink, had very high validation performance, followed by the second most-numerous class, niche shift (Fig. 2). Species in the “stable” class were often misclassified as shifts. The main validation problem though occurred for species undergoing niche expansion, the least numerous class (representing just 8% of species), which were typically misclassified. This outcome is in part related to the level of tolerance of 20%, which does not classify niche increases as expansions (or niche contractions as shrinks) unless these are substantial. In fact, with a tolerance level of 5% there would be more than twice as many species classified as niche expansions, and the model is slightly better able at classifying them (but much less able at classifying shifting species).

**Figure 2.**
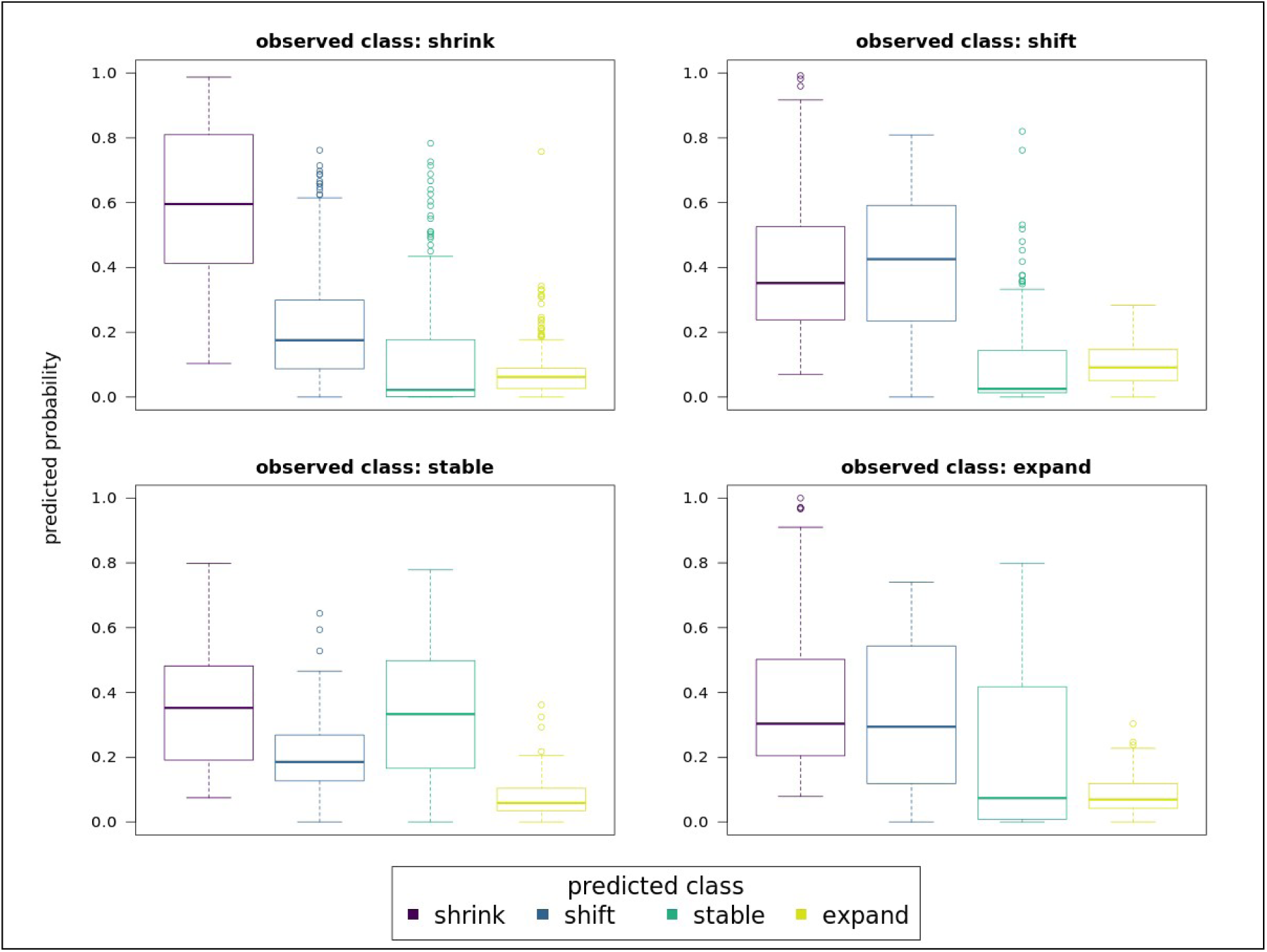
Probability to belong to different classes of niche change, for species in different observed categories, as predicted by the model based on Mid Holocene climate (under global circulation model IPSL-CM5A-LR) and a tolerance of 20% to separate niche change from niche stability. Each set of boxplots reports the probability of species within an observed niche category (reported in the plot title) to belong to any of the 4 categories.

We did not use phylogenetic relationships as predictors in our final model, after verifying that including phylogenetic eigenvectors did not improve the AIC of our best performing model based on IPSL-CM5A-LR MID climate and a tolerance of 20% (Table S4). The two imputed variables in our model, gestation length and inter-birth interval, had acceptable performances during imputation, in terms of normalised root mean square error (NRMSE), with gestation length (NRMSE = 0.22) performing better than inter-birth interval (NRMSE = 0.32). We only used one imputation (out of ten imputations run), as the effect of phylogenetic uncertainty on data imputation was negligible when predicting species niche classes. In fact, for 98% of the species the predicted class (i.e. the one with the highest probability from the multinomial model) did not change when using intrinsic traits imputed from 10 alternative phylogenies (Table S5).

### Drivers of change in species climatic niches

After averaging the result of niche predictions based on the two GCMs for the MID climate, we found that different niche categories were best predicted by different sets of variables (Fig. 3). As expected, we found overall climatic stability was a strong discriminant of stable *vs* shrinking niche, with a less strong effect on the prediction of other classes (Fig. 3, Table S6). Invertebrate diet and current land use are relatively strong predictors of niche shift, together with body mass and interbirth interval. These latter two variables are also relatively important predictor of niche expansion. Past human population density and current land-use are relatively important predictors of niche stability and shift, respectively. Variable importance measured in a random forest model for classification reflected the overall patterns of the multinomial model, with the most important variables being climatic stability, biogeographic realm, past human population, current land-use, and body mass (Fig. S1).

**Figure 3.**
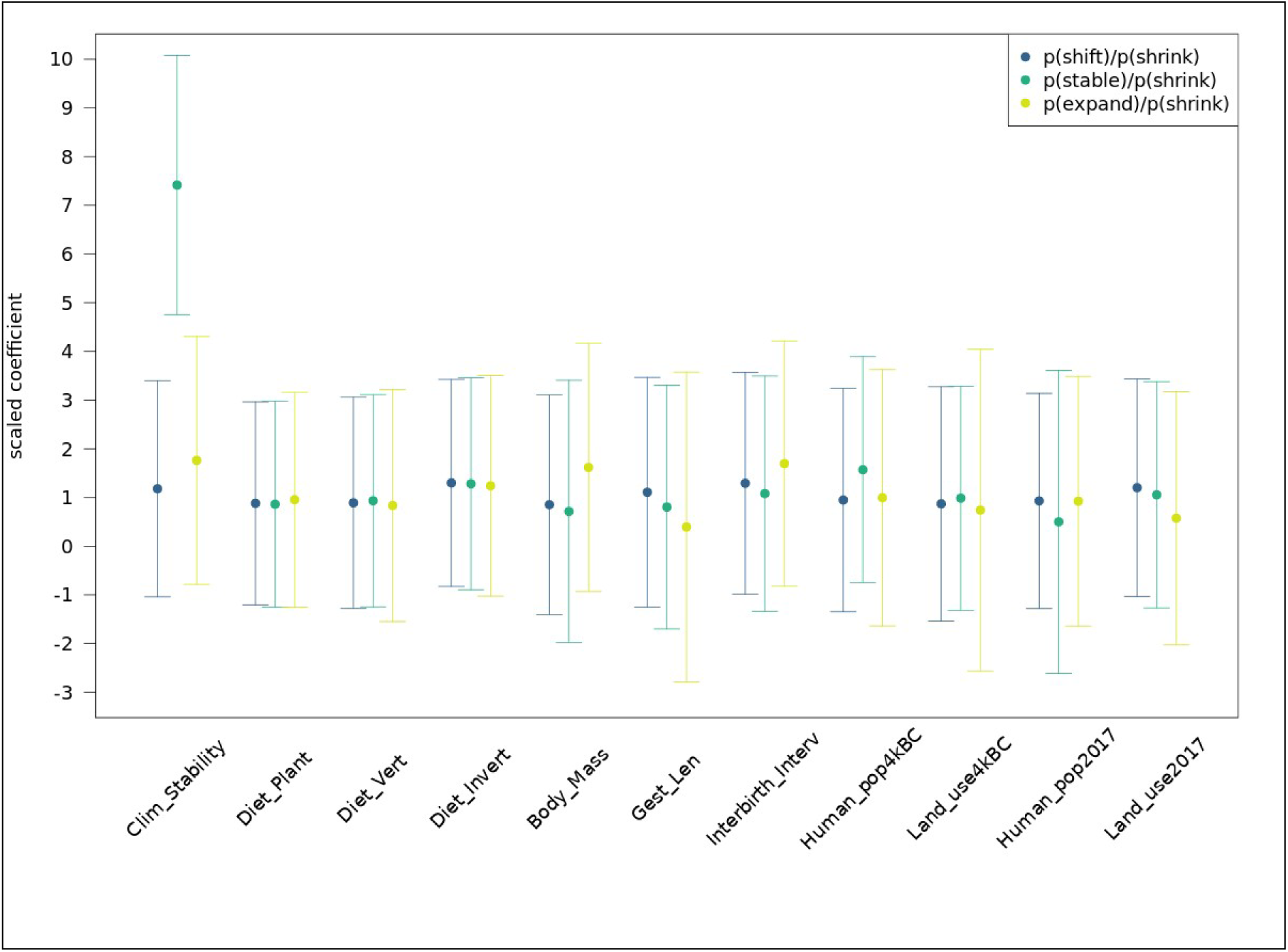
Averaged model’s coefficients for the prediction of species niche change categories, using Mid Holocene as the reference past climate and two alternative GCMs. For each quantitative predictor variable used in the multinomial models, the scaled coefficient is reported representing the odds of being in the categories “shift”, “stable”, or “expand” rather than the class “shrink” (used as a reference).

When considering diet, we found species consuming higher percentage of invertebrate food had a lower probability of undergoing niche shrink and higher probability of niche expansion or shift (Fig. 4a,c,e). Species consuming more vertebrate or plant food showed the opposite pattern. When considering life-history traits instead, we found species were more likely to undergo niche shrink as their body mass increases, with the exception of very large species (pachyderms) which are less likely to undergo niche shrink (Fig. 4b). Larger species were also much more likely to expand their niche, and less likely to show niche shift or stability. Species with longer gestation time – i.e. low reproductive output, *sensu* Bielby et al. (2007) - were much more likely to undergo niche shrink and less likely to belong to any other class of niche change (Fig. 5d). Species with longer interbirth intervals - slow reproductive timing, *sensu* Bielby et al. (2007) - instead were less likely to have a shrink or stable niche, but more likely to shift their climatic niche (Fig. 4f). This same pattern was observed when replacing interbirth interval with other variables representing reproductive timing, such as weaning age or age at sexual maturity (Fig. S2). However these variables had even lower performance during data imputation routine (NRMSE was 0.42 and 0.36 respectively).

**Figure 4.**
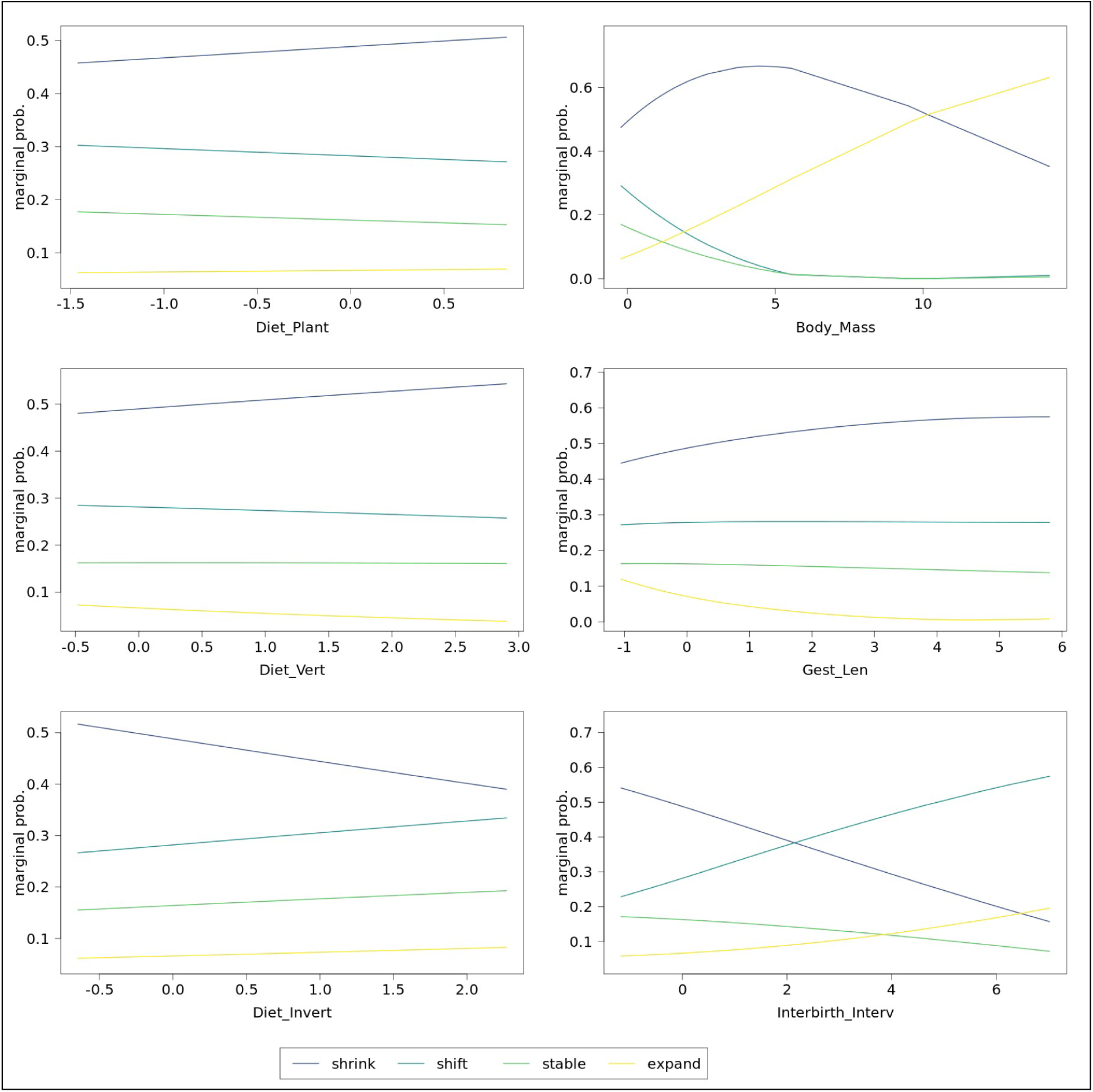
Average partial effect plots of the relationship between species intrinsic characteristics (scaled for the multinomial model) and probability of species to be assigned to one of four categories of niche change (shrink, stable, shift, expand), using Mid Holocene as the reference past climate and two alternative GCMs. Panels a,c,e represents species diets as proportion of plant, vertebrate, and invertebrate food consumed. Panels b,d,f represents species’ body mass, gestation length, and inter-birth interval.

**Figure 5.**
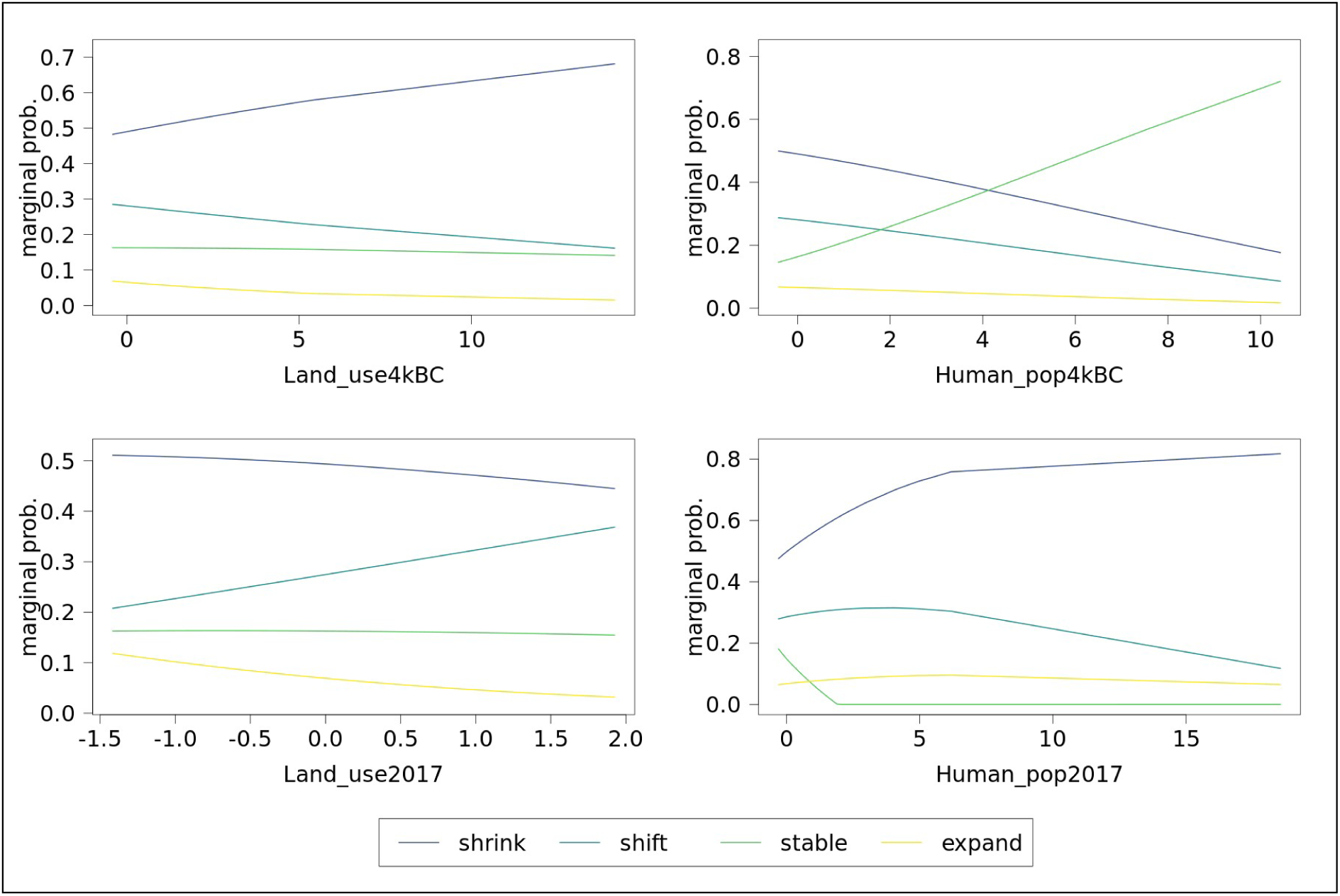
Average partial effect plot of the relationship between human pressure (scaled for the multinomial model) and probability of species to be assigned to one of four categories of niche change (shrink, stable, shift, expand), using Mid Holocene as the reference past climate and two alternative GCMs. Panels a,c represents historic and current human land use, panels b,d represents historic and current human population.

When looking at human pressure, we found past and current variables have somewhat different relationships with niche class probabilities (Fig. 5). Higher probability of niche shrink is associated with past agricultural land-use, while current land-use is associated with higher probability of niche shift. Human population density instead shows different patterns, as past density is associated with higher probability of niche stability and current density is associated with higher probability of niche shrink.

## Discussion

Overall, half of the species we analysed have faced a shrinkage in their realised climatic niche of more than 20%, in response to human alteration of their distribution, global climatic change, and life history traits. At the same time, only 15-18% of species retained a stable or nearly stable niche. We used 20% as a threshold to separate stability from change, both in terms of change in niche position and change in overall niche variability. This is a quite conservative threshold, which was chosen for practical and statistical reasons. Practically, given the coarse resolution of past species distribution data, choosing a relatively high threshold minimised the risk of identifying changes that were an artefact of data uncertainty. Statistically, this choice resulted in the best performance of the model, which was higher with threshold of 20% especially when looking at category-level classification accuracy.

We found species with certain biological characteristics were more likely to undergo niche shrink. Large-bodied species for example were more likely to undergo niche shrink compared to smaller species, and less likely to show niche shift or stability. There was an exception however for very large species, i.e. pachyderms, which might be due to the large conservation attention that some of these species receive nowadays. A correlation between niche shrink and large body mass might depend on the large mammals’ vulnerability to human impact, which determines low resistance to niche erosion. Larger species have the biological potential to extend their distribution range via dispersal mechanisms (Santini et al. 2013), but are also typically characterised by slower life histories compared to smaller species (Bielby et al. 2007) and are more sensitive to human impact (Cardillo et al. 2005). In fact, this result is reflected when looking at gestation length, a proxy of reproductive output (Bielby et al. 2007), with longer values associated with higher probability of niche shrinkage. Instead interbirth interval, a proxy of reproductive timing (Bielby et al. 2007), was positively associated with niche shift and negatively associated with niche shrink. However there was limited available data on this trait (and other reproductive timing traits), and the data imputation procedure had lower performance for this variable compared to gestation length. Improved information would be needed before drawing conclusions on this variable. Species’ diet was not an overall dominant driver of niche change in our model, but showed fairly clear relationship with niche categories. We found the main effect was determined by invertebrate food items, with higher percentage of invertebrate diet leading to lower probability of niche shrink and higher probability of expansion. This might imply highly insectivorous species have higher adaptability compared to both highly carnivorous and highly herbivorous ones, similar to omnivores.

We found species responded to past and current levels of human pressure in different ways. Higher levels of historical land-use change within a species’ natural distribution determined higher probability of niche shrink. Current levels of land-use change instead determined higher probability of shift. This result might have emerged because part of the current human influence is realized over portions of the natural species distributions which have been lost. Species might be able to adapt to human pressure inside their present-day distributions, which already resisted to some level of historical pressure. When low levels of climate change affect these core distribution areas, species might show some adaptation capacity via niche shift mechanisms. The relationship with human population density showed the opposite pattern instead, with past density positively correlated with the probability of stable niche and negatively correlated with the probability of shrinking niche. This scenario of past co-existence between human and other mammal species might be a reflection of both human and animal communities settling in productive natural environments, but also a possible facilitation of human-wildlife coexistence in those areas when human colonization started earlier (Carter & Linnell 2016). This result however is also in part dependent on the threshold of 20% that was used to distinguish between niche stability and niche change. In fact, we verified that a lower threshold of 5% would result in a positive correlation between past human density and probability of niche shrink; this means past human density resulted in either niche stability or very low levels of niche shrink. Instead we found current human population density within natural species distributions was positively associated with niche shrink, regardless of the threshold. This shows the risk faced by species living in highly anthropogenic areas (Di Marco et al. 2018), which might become more vulnerable to the additive effect of climate change (Mantyka-Pringle et al. 2012, 2015).

Our model has demonstrated good overall performance during validation, except for the ‘expand’ category. This is due to the limited number of species which showed a substantial (>20%) increase in their niche breadth over time. This implies our understanding of the mechanisms of niche expansion is still limited, until additional species examples are identified. A promising field of research in this case is represented by invasive species (Broennimann et al. 2012). Invasive species might maintain their original climatic niche in the invaded region (Petitpierre et al. 2012), or exploit a wider variety of climatic conditions and shift or expand their realised niche (Lauzeral et al. 2011). Understanding more of the dynamics and drivers of niche change for these species can shed light on the past dynamics of niche change.

We identified the conditions under which species are unlikely to maintain a varied climatic niche, potentially losing their climate adaptive potential. Areas which will experience substantially different climates in the future should be given special attention to prevent threatened and restricted-range species from rapid decline. Interventions that facilitate natural dispersal, or assisted colonisation, should be carefully evaluated for these species, as part of international strategies to combat the effects of climate change on biodiversity such as the Convention on Biological Diversity and the United Nation Framework Convention on Climate Change.

## Acknowledgements

This project has received funding from the European Union’s Horizon 2020 research and innovation programme under the Marie Skłodowska-Curie grant agreement No. 793212.

