## Supplementary material for "Drivers of change in the realised climatic niche of terrestrial mammal species": Table S1

***Table S1*** *List of species analysed (and their taxonomy).*

| Binomial | Order | Family |
|---|---|---|
| Echinops telfairi | Afrosoricida | Tenrecidae |
| Geogale aurita | Afrosoricida | Tenrecidae |
| Hemicentetes nigriceps | Afrosoricida | Tenrecidae |
| Hemicentetes semispinosus | Afrosoricida | Tenrecidae |
| Limnogale mergulus | Afrosoricida | Tenrecidae |
| Microgale brevicaudata | Afrosoricida | Tenrecidae |
| Microgale cowani | Afrosoricida | Tenrecidae |
| Microgale dobsoni | Afrosoricida | Tenrecidae |
| Microgale drouhardi | Afrosoricida | Tenrecidae |
| Microgale dryas | Afrosoricida | Tenrecidae |
| Microgale fotsifotsy | Afrosoricida | Tenrecidae |
| Microgale gracilis | Afrosoricida | Tenrecidae |
| Microgale grandidieri | Afrosoricida | Tenrecidae |
| Microgale gymnorhyncha | Afrosoricida | Tenrecidae |
| Microgale majori | Afrosoricida | Tenrecidae |
| Microgale monticola | Afrosoricida | Tenrecidae |
| Microgale nasoloi | Afrosoricida | Tenrecidae |
| Microgale parvula | Afrosoricida | Tenrecidae |
| Microgale principula | Afrosoricida | Tenrecidae |
| Microgale pusilla | Afrosoricida | Tenrecidae |
| Microgale soricoides | Afrosoricida | Tenrecidae |
| Microgale taiva | Afrosoricida | Tenrecidae |
| Microgale talazaci | Afrosoricida | Tenrecidae |
| Microgale thomasi | Afrosoricida | Tenrecidae |
| Oryzorictes hova | Afrosoricida | Tenrecidae |
| Ailurus fulgens | Carnivora | Ailuridae |
| Canis latrans | Carnivora | Canidae |
| Canis lupus | Carnivora | Canidae |
| Canis mesomelas | Carnivora | Canidae |
| Canis rufus | Carnivora | Canidae |
| Canis simensis | Carnivora | Canidae |
| Cuon alpinus | Carnivora | Canidae |
| Dusicyon australis | Carnivora | Canidae |
| Lycaon pictus | Carnivora | Canidae |
| Vulpes velox | Carnivora | Canidae |
| Vulpes vulpes | Carnivora | Canidae |
| Cryptoprocta ferox | Carnivora | Eupleridae |
| Cryptoprocta spelea | Carnivora | Eupleridae |
| Eupleres goudotii | Carnivora | Eupleridae |
| Eupleres major | Carnivora | Eupleridae |
| Fossa fossana | Carnivora | Eupleridae |
| Galidia elegans | Carnivora | Eupleridae |
| Galidictis fasciata | Carnivora | Eupleridae |
| Mungotictis decemlineata | Carnivora | Eupleridae |
| Salanoia concolor | Carnivora | Eupleridae |

| | | |
|---|---|---|
| Acinonyx jubatus | Carnivora | Felidae |
| Caracal aurata | Carnivora | Felidae |
| Caracal caracal | Carnivora | Felidae |
| Catopuma badia | Carnivora | Felidae |
| Catopuma temminckii | Carnivora | Felidae |
| Felis chaus | Carnivora | Felidae |
| Felis margarita | Carnivora | Felidae |
| Felis silvestris | Carnivora | Felidae |
| Herpailurus yagouaroundi | Carnivora | Felidae |
| Leopardus pardalis | Carnivora | Felidae |
| Leptailurus serval | Carnivora | Felidae |
| Lynx canadensis | Carnivora | Felidae |
| Lynx lynx | Carnivora | Felidae |
| Lynx pardinus | Carnivora | Felidae |
| Lynx rufus | Carnivora | Felidae |
| Neofelis diardi | Carnivora | Felidae |
| Neofelis nebulosa | Carnivora | Felidae |
| Panthera leo | Carnivora | Felidae |
| Panthera onca | Carnivora | Felidae |
| Panthera pardus | Carnivora | Felidae |
| Panthera tigris | Carnivora | Felidae |
| Pardofelis marmorata | Carnivora | Felidae |
| Prionailurus viverrinus | Carnivora | Felidae |
| Puma concolor | Carnivora | Felidae |
| Crocuta crocuta | Carnivora | Hyaenidae |
| Hyaena hyaena | Carnivora | Hyaenidae |
| Parahyaena brunnea | Carnivora | Hyaenidae |
| Proteles cristata | Carnivora | Hyaenidae |
| Enhydra lutris | Carnivora | Mustelidae |
| Gulo gulo | Carnivora | Mustelidae |
| Lontra canadensis | Carnivora | Mustelidae |
| Lutra lutra | Carnivora | Mustelidae |
| Lutra sumatrana | Carnivora | Mustelidae |
| Martes americana | Carnivora | Mustelidae |
| Martes gwatkinsii | Carnivora | Mustelidae |
| Martes martes | Carnivora | Mustelidae |
| Martes melampus | Carnivora | Mustelidae |
| Martes pennanti | Carnivora | Mustelidae |
| Martes zibellina | Carnivora | Mustelidae |
| Melogale everetti | Carnivora | Mustelidae |
| Mustela lutreola | Carnivora | Mustelidae |
| Mustela lutreolina | Carnivora | Mustelidae |
| Mustela nivalis | Carnivora | Mustelidae |
| Neovison macrodon | Carnivora | Mustelidae |
| Neovison vison | Carnivora | Mustelidae |
| Phocarctos hookeri | Carnivora | Otariidae |
| Zalophus japonicus | Carnivora | Otariidae |
| Monachus monachus | Carnivora | Phocidae |
| Neomonachus tropicalis | Carnivora | Phocidae |

| | | |
|---|---|---|
| Pagophilus groenlandicus | Carnivora | Phocidae |
| Prionodon linsang | Carnivora | Prionodontidae |
| Ailuropoda melanoleuca | Carnivora | Ursidae |
| Tremarctos ornatus | Carnivora | Ursidae |
| Ursus americanus | Carnivora | Ursidae |
| Ursus arctos | Carnivora | Ursidae |
| Genetta piscivora | Carnivora | Viverridae |
| Macrogalidia musschenbroekii | Carnivora | Viverridae |
| Poiana leightoni | Carnivora | Viverridae |
| Viverra megaspila | Carnivora | Viverridae |
| Antilocapra americana | Cetartiodactyla | Antilocapridae |
| Addax nasomaculatus | Cetartiodactyla | Bovidae |
| Alcelaphus buselaphus | Cetartiodactyla | Bovidae |
| Ammodorcas clarkei | Cetartiodactyla | Bovidae |
| Ammotragus lervia | Cetartiodactyla | Bovidae |
| Antilope cervicapra | Cetartiodactyla | Bovidae |
| Beatragus hunteri | Cetartiodactyla | Bovidae |
| Bison bison | Cetartiodactyla | Bovidae |
| Bison bonasus | Cetartiodactyla | Bovidae |
| Bos gaurus | Cetartiodactyla | Bovidae |
| Bos javanicus | Cetartiodactyla | Bovidae |
| Bos mutus | Cetartiodactyla | Bovidae |
| Bos primigenius | Cetartiodactyla | Bovidae |
| Boselaphus tragocamelus | Cetartiodactyla | Bovidae |
| Bubalus arnee | Cetartiodactyla | Bovidae |
| Bubalus depressicornis | Cetartiodactyla | Bovidae |
| Bubalus mindorensis | Cetartiodactyla | Bovidae |
| Capra aegagrus | Cetartiodactyla | Bovidae |
| Capra falconeri | Cetartiodactyla | Bovidae |
| Capra ibex | Cetartiodactyla | Bovidae |
| Capra nubiana | Cetartiodactyla | Bovidae |
| Capra pyrenaica | Cetartiodactyla | Bovidae |
| Capricornis swinhoei | Cetartiodactyla | Bovidae |
| Cephalophus zebra | Cetartiodactyla | Bovidae |
| Damaliscus lunatus | Cetartiodactyla | Bovidae |
| Eudorcas rufifrons | Cetartiodactyla | Bovidae |
| Gazella bilkis | Cetartiodactyla | Bovidae |
| Gazella dorcas | Cetartiodactyla | Bovidae |
| Gazella gazella | Cetartiodactyla | Bovidae |
| Gazella saudiya | Cetartiodactyla | Bovidae |
| Hippotragus equinus | Cetartiodactyla | Bovidae |
| Hippotragus leucophaeus | Cetartiodactyla | Bovidae |
| Naemorhedus caudatus | Cetartiodactyla | Bovidae |
| Nanger dama | Cetartiodactyla | Bovidae |
| Nanger soemmerringii | Cetartiodactyla | Bovidae |
| Nilgiritragus hylocrius | Cetartiodactyla | Bovidae |
| Oreamnos americanus | Cetartiodactyla | Bovidae |
| Oryx beisa | Cetartiodactyla | Bovidae |
| Oryx dammah | Cetartiodactyla | Bovidae |

| | | |
|---|---|---|
| Oryx leucoryx | Cetartiodactyla | Bovidae |
| Ovibos moschatus | Cetartiodactyla | Bovidae |
| Ovis ammon | Cetartiodactyla | Bovidae |
| Ovis canadensis | Cetartiodactyla | Bovidae |
| Ovis orientalis | Cetartiodactyla | Bovidae |
| Pantholops hodgsonii | Cetartiodactyla | Bovidae |
| Procapra przewalskii | Cetartiodactyla | Bovidae |
| Pseudoryx nghetinhensis | Cetartiodactyla | Bovidae |
| Raphicerus campestris | Cetartiodactyla | Bovidae |
| Rupicapra pyrenaica | Cetartiodactyla | Bovidae |
| Saiga tatarica | Cetartiodactyla | Bovidae |
| Syncerus caffer | Cetartiodactyla | Bovidae |
| Tragelaphus derbianus | Cetartiodactyla | Bovidae |
| Camelus ferus | Cetartiodactyla | Camelidae |
| Alces alces | Cetartiodactyla | Cervidae |
| Axis axis | Cetartiodactyla | Cervidae |
| Axis kuhlii | Cetartiodactyla | Cervidae |
| Axis porcinus | Cetartiodactyla | Cervidae |
| Cervus canadensis | Cetartiodactyla | Cervidae |
| Cervus elaphus | Cetartiodactyla | Cervidae |
| Cervus nippon | Cetartiodactyla | Cervidae |
| Dama dama | Cetartiodactyla | Cervidae |
| Dama mesopotamica | Cetartiodactyla | Cervidae |
| Elaphurus davidianus | Cetartiodactyla | Cervidae |
| Hippocamelus antisensis | Cetartiodactyla | Cervidae |
| Hippocamelus bisulcus | Cetartiodactyla | Cervidae |
| Hydropotes inermis | Cetartiodactyla | Cervidae |
| Muntiacus crinifrons | Cetartiodactyla | Cervidae |
| Odocoileus hemionus | Cetartiodactyla | Cervidae |
| Ozotoceros bezoarticus | Cetartiodactyla | Cervidae |
| Rangifer tarandus | Cetartiodactyla | Cervidae |
| Rucervus duvaucelii | Cetartiodactyla | Cervidae |
| Rucervus eldii | Cetartiodactyla | Cervidae |
| Rucervus schomburgki | Cetartiodactyla | Cervidae |
| Rusa alfredi | Cetartiodactyla | Cervidae |
| Rusa marianna | Cetartiodactyla | Cervidae |
| Rusa timorensis | Cetartiodactyla | Cervidae |
| Giraffa camelopardalis | Cetartiodactyla | Giraffidae |
| Hippopotamus amphibius | Cetartiodactyla | Hippopotamidae |
| Hippopotamus lemerlei | Cetartiodactyla | Hippopotamidae |
| Hippopotamus madagascariensis | Cetartiodactyla | Hippopotamidae |
| Porcula salvania | Cetartiodactyla | Suidae |
| Sus barbatus | Cetartiodactyla | Suidae |
| Sus bucculentus | Cetartiodactyla | Suidae |
| Sus cebifrons | Cetartiodactyla | Suidae |
| Sus oliveri | Cetartiodactyla | Suidae |
| Sus philippensis | Cetartiodactyla | Suidae |
| Sus scrofa | Cetartiodactyla | Suidae |

| | | |
|---|---|---|
| Sus verrucosus | Cetartiodactyla | Suidae |
| Catagonus wagneri | Cetartiodactyla | Tayassuidae |
| Taphozous hildegardeae | Chiroptera | Emballonuridae |
| Hipposideros grandis | Chiroptera | Hipposideridae |
| Hipposideros inornatus | Chiroptera | Hipposideridae |
| Hipposideros orbiculus | Chiroptera | Hipposideridae |
| Hipposideros ridleyi | Chiroptera | Hipposideridae |
| Macroderma gigas | Chiroptera | Megadermatidae |
| Chaerephon bregullae | Chiroptera | Molossidae |
| Chaerephon johorensis | Chiroptera | Molossidae |
| Eumops floridanus | Chiroptera | Molossidae |
| Mormopterus acetabulosus | Chiroptera | Molossidae |
| Mormopterus minutus | Chiroptera | Molossidae |
| Mystacina robusta | Chiroptera | Mystacinidae |
| Mystacina tuberculata | Chiroptera | Mystacinidae |
| Natalus primus | Chiroptera | Natalidae |
| Nycteris javanica | Chiroptera | Nycteridae |
| Vampyressa melissa | Chiroptera | Phyllostomidae |
| Acerodon jubatus | Chiroptera | Pteropodidae |
| Aproteles bulmerae | Chiroptera | Pteropodidae |
| Dobsonia chapmani | Chiroptera | Pteropodidae |
| Dobsonia emersa | Chiroptera | Pteropodidae |
| Megaerops wetmorei | Chiroptera | Pteropodidae |
| Mirimiri acrodonta | Chiroptera | Pteropodidae |
| Pteralopex anceps | Chiroptera | Pteropodidae |
| Pteropus fundatus | Chiroptera | Pteropodidae |
| Pteropus pilosus | Chiroptera | Pteropodidae |
| Pteropus poliocephalus | Chiroptera | Pteropodidae |
| Pteropus rodricensis | Chiroptera | Pteropodidae |
| Pteropus subniger | Chiroptera | Pteropodidae |
| Pteropus tokudae | Chiroptera | Pteropodidae |
| Rhinolophus maclaudi | Chiroptera | Rhinolophidae |
| Rhinolophus mehelyi | Chiroptera | Rhinolophidae |
| Rhinolophus montanus | Chiroptera | Rhinolophidae |
| Chalinolobus tuberculatus | Chiroptera | Vespertilionidae |
| Eptesicus diminutus | Chiroptera | Vespertilionidae |
| Eptesicus japonensis | Chiroptera | Vespertilionidae |
| Hesperoptenus tomesi | Chiroptera | Vespertilionidae |
| Murina ryukyuana | Chiroptera | Vespertilionidae |
| Murina tenebrosa | Chiroptera | Vespertilionidae |
| Myotis capaccinii | Chiroptera | Vespertilionidae |
| Myotis pruinosus | Chiroptera | Vespertilionidae |
| Myotis vivesi | Chiroptera | Vespertilionidae |
| Myotis yanbarensis | Chiroptera | Vespertilionidae |
| Nyctalus furvus | Chiroptera | Vespertilionidae |
| Nyctophilus nebulosus | Chiroptera | Vespertilionidae |
| Pipistrellus endoi | Chiroptera | Vespertilionidae |
| Dasypus novemcinctus | Cingulata | CingulataFam |
| Dasycercus blythi | Dasyuromorphia | Dasyuridae |

| | | |
|---|---|---|
| Dasycercus cristicauda | Dasyuromorphia | Dasyuridae |
| Dasyuroides byrnei | Dasyuromorphia | Dasyuridae |
| Dasyurus geoffroii | Dasyuromorphia | Dasyuridae |
| Dasyurus hallucatus | Dasyuromorphia | Dasyuridae |
| Dasyurus maculatus | Dasyuromorphia | Dasyuridae |
| Parantechinus apicalis | Dasyuromorphia | Dasyuridae |
| Phascogale calura | Dasyuromorphia | Dasyuridae |
| Phascogale pirata | Dasyuromorphia | Dasyuridae |
| Phascogale tapoatafa | Dasyuromorphia | Dasyuridae |
| Pseudantechinus mimulus | Dasyuromorphia | Dasyuridae |
| Sarcophilus harrisii | Dasyuromorphia | Dasyuridae |
| Sminthopsis gilberti | Dasyuromorphia | Dasyuridae |
| Sminthopsis psammophila | Dasyuromorphia | Dasyuridae |
| Myrmecobius fasciatus | Dasyuromorphia | Myrmecobiidae |
| Thylacinus cynocephalus | Dasyuromorphia | Thylacinidae |
| Chacodelphys formosa | Didelphimorphia | Didelphidae |
| Cryptonanus ignitus | Didelphimorphia | Didelphidae |
| Marmosops handleyi | Didelphimorphia | Didelphidae |
| Acrobates pygmaeus | Diprotodontia | Acrobatidae |
| Dendrolagus goodfellowi | Diprotodontia | Macropodidae |
| Dendrolagus inustus | Diprotodontia | Macropodidae |
| Dendrolagus matschiei | Diprotodontia | Macropodidae |
| Dendrolagus spadix | Diprotodontia | Macropodidae |
| Dendrolagus ursinus | Diprotodontia | Macropodidae |
| Dorcopsis luctuosa | Diprotodontia | Macropodidae |
| Lagorchestes asomatus | Diprotodontia | Macropodidae |
| Lagorchestes hirsutus | Diprotodontia | Macropodidae |
| Lagorchestes leporides | Diprotodontia | Macropodidae |
| Lagostrophus fasciatus | Diprotodontia | Macropodidae |
| Macropus eugenii | Diprotodontia | Macropodidae |
| Macropus greyi | Diprotodontia | Macropodidae |
| Macropus parryi | Diprotodontia | Macropodidae |
| Onychogalea fraenata | Diprotodontia | Macropodidae |
| Onychogalea lunata | Diprotodontia | Macropodidae |
| Petrogale lateralis | Diprotodontia | Macropodidae |
| Petrogale xanthopus | Diprotodontia | Macropodidae |
| Setonix brachyurus | Diprotodontia | Macropodidae |
| Thylogale billardierii | Diprotodontia | Macropodidae |
| Thylogale brunii | Diprotodontia | Macropodidae |
| Thylogale calabyi | Diprotodontia | Macropodidae |
| Gymnobelideus leadbeateri | Diprotodontia | Petauridae |
| Petaurus australis | Diprotodontia | Petauridae |
| Trichosurus vulpecula | Diprotodontia | Phalangeridae |
| Phascolarctos cinereus | Diprotodontia | Phascolarctidae |
| Aepyprymnus rufescens | Diprotodontia | Potoroidae |
| Bettongia anhydra | Diprotodontia | Potoroidae |
| Bettongia gaimardi | Diprotodontia | Potoroidae |
| Bettongia lesueur | Diprotodontia | Potoroidae |
| Bettongia penicillata | Diprotodontia | Potoroidae |

| | | |
|---|---|---|
| Bettongia pusilla | Diprotodontia | Potoroidae |
| Bettongia tropica | Diprotodontia | Potoroidae |
| Caloprymnus campestris | Diprotodontia | Potoroidae |
| Potorous gilbertii | Diprotodontia | Potoroidae |
| Potorous platyops | Diprotodontia | Potoroidae |
| Pseudocheirus occidentalis | Diprotodontia | Pseudocheiridae |
| Pseudocheirus peregrinus | Diprotodontia | Pseudocheiridae |
| Pseudochirulus schlegeli | Diprotodontia | Pseudocheiridae |
| Lasiorhinus krefftii | Diprotodontia | Vombatidae |
| Lasiorhinus latifrons | Diprotodontia | Vombatidae |
| Neohylomys hainanensis | Eulipotyphla | Erinaceidae |
| Nesophontes edithae | Eulipotyphla | Nesophontidae |
| Nesophontes hypomicrus | Eulipotyphla | Nesophontidae |
| Nesophontes major | Eulipotyphla | Nesophontidae |
| Nesophontes micrus | Eulipotyphla | Nesophontidae |
| Nesophontes paramicrus | Eulipotyphla | Nesophontidae |
| Nesophontes zamicrus | Eulipotyphla | Nesophontidae |
| Solenodon cubanus | Eulipotyphla | Solenodontidae |
| Solenodon marcanoi | Eulipotyphla | Solenodontidae |
| Solenodon paradoxus | Eulipotyphla | Solenodontidae |
| Crocidura andamanensis | Eulipotyphla | Soricidae |
| Crocidura baileyi | Eulipotyphla | Soricidae |
| Crocidura desperata | Eulipotyphla | Soricidae |
| Crocidura hispida | Eulipotyphla | Soricidae |
| Crocidura jenkinsi | Eulipotyphla | Soricidae |
| Crocidura miya | Eulipotyphla | Soricidae |
| Crocidura pachyura | Eulipotyphla | Soricidae |
| Crocidura stenocephala | Eulipotyphla | Soricidae |
| Crocidura suaveolens | Eulipotyphla | Soricidae |
| Crocidura tarella | Eulipotyphla | Soricidae |
| Crocidura telfordi | Eulipotyphla | Soricidae |
| Crocidura virgata | Eulipotyphla | Soricidae |
| Cryptotis obscura | Eulipotyphla | Soricidae |
| Sorex sclateri | Eulipotyphla | Soricidae |
| Suncus mertensi | Eulipotyphla | Soricidae |
| Sylvisorex camerunensis | Eulipotyphla | Soricidae |
| Desmana moschata | Eulipotyphla | Talpidae |
| Lepus corsicanus | Lagomorpha | Leporidae |
| Lepus europaeus | Lagomorpha | Leporidae |
| Lepus flavigularis | Lagomorpha | Leporidae |
| Oryctolagus cuniculus | Lagomorpha | Leporidae |
| Sylvilagus transitionalis | Lagomorpha | Leporidae |
| Ochotona hoffmanni | Lagomorpha | Ochotonidae |
| Prolagus sardus | Lagomorpha | Prolagidae |
| Rhynchocyon chrysopygus | Macroscelidea | Macroscelididae |
| Zaglossus bartoni | Monotremata | Tachyglossidae |
| Zaglossus bruijnii | Monotremata | Tachyglossidae |
| Caenolestes convelatus | Paucituberculata | Caenolestidae |
| Caenolestes sangay | Paucituberculata | Caenolestidae |

| | | |
|---|---|---|
| Chaeropus ecaudatus | Peramelemorphia | Chaeropodidae |
| Isoodon auratus | Peramelemorphia | Peramelidae |
| Isoodon obesulus | Peramelemorphia | Peramelidae |
| Perameles bougainville | Peramelemorphia | Peramelidae |
| Perameles eremiana | Peramelemorphia | Peramelidae |
| Perameles gunnii | Peramelemorphia | Peramelidae |
| Rhynchomeles prattorum | Peramelemorphia | Peramelidae |
| Macrotis lagotis | Peramelemorphia | Thylacomyidae |
| Macrotis leucura | Peramelemorphia | Thylacomyidae |
| Equus africanus | Perissodactyla | Equidae |
| Equus ferus | Perissodactyla | Equidae |
| Equus grevyi | Perissodactyla | Equidae |
| Equus hemionus | Perissodactyla | Equidae |
| Equus quagga | Perissodactyla | Equidae |
| Equus zebra | Perissodactyla | Equidae |
| Ceratotherium simum | Perissodactyla | Rhinocerotidae |
| Dicerorhinus sumatrensis | Perissodactyla | Rhinocerotidae |
| Diceros bicornis | Perissodactyla | Rhinocerotidae |
| Rhinoceros sondaicus | Perissodactyla | Rhinocerotidae |
| Rhinoceros unicornis | Perissodactyla | Rhinocerotidae |
| Tapirus indicus | Perissodactyla | Tapiridae |
| Tapirus pinchaque | Perissodactyla | Tapiridae |
| Manis javanica | Pholidota | Manidae |
| Phataginus tetradactyla | Pholidota | Manidae |
| Smutsia gigantea | Pholidota | Manidae |
| Bradypus torquatus | Pilosa | Bradypodidae |
| Ateles hybridus | Primates | Atelidae |
| Brachyteles hypoxanthus | Primates | Atelidae |
| Allochrocebus preussi | Primates | Cercopithecidae |
| Macaca arctoides | Primates | Cercopithecidae |
| Macaca cyclopis | Primates | Cercopithecidae |
| Macaca sinica | Primates | Cercopithecidae |
| Macaca sylvanus | Primates | Cercopithecidae |
| Mandrillus leucophaeus | Primates | Cercopithecidae |
| Nasalis larvatus | Primates | Cercopithecidae |
| Piliocolobus badius | Primates | Cercopithecidae |
| Piliocolobus tephrosceles | Primates | Cercopithecidae |
| Presbytis chrysomelas | Primates | Cercopithecidae |
| Rhinopithecus avunculus | Primates | Cercopithecidae |
| Rhinopithecus bieti | Primates | Cercopithecidae |
| Rhinopithecus roxellana | Primates | Cercopithecidae |
| Rhinopithecus strykeri | Primates | Cercopithecidae |
| Rungwecebus kipunji | Primates | Cercopithecidae |
| Trachypithecus francoisi | Primates | Cercopithecidae |
| Trachypithecus vetulus | Primates | Cercopithecidae |
| Allocebus trichotis | Primates | Cheirogaleidae |
| Cheirogaleus medius | Primates | Cheirogaleidae |
| Microcebus bongolavensis | Primates | Cheirogaleidae |
| Microcebus griseorufus | Primates | Cheirogaleidae |

| | | |
|---|---|---|
| Microcebus jollyae | Primates | Cheirogaleidae |
| Microcebus murinus | Primates | Cheirogaleidae |
| Microcebus ravelobensis | Primates | Cheirogaleidae |
| Microcebus rufus | Primates | Cheirogaleidae |
| Microcebus tavaratra | Primates | Cheirogaleidae |
| Mirza coquereli | Primates | Cheirogaleidae |
| Mirza zaza | Primates | Cheirogaleidae |
| Phaner electromontis | Primates | Cheirogaleidae |
| Phaner furcifer | Primates | Cheirogaleidae |
| Phaner pallescens | Primates | Cheirogaleidae |
| Phaner parienti | Primates | Cheirogaleidae |
| Daubentonia madagascariensis | Primates | Daubentoniidae |
| Galagoides rondoensis | Primates | Galagidae |
| Gorilla gorilla | Primates | Hominidae |
| Pan troglodytes | Primates | Hominidae |
| Pongo abelii | Primates | Hominidae |
| Pongo pygmaeus | Primates | Hominidae |
| Hoolock hoolock | Primates | Hylobatidae |
| Nomascus concolor | Primates | Hylobatidae |
| Nomascus gabriellae | Primates | Hylobatidae |
| Nomascus hainanus | Primates | Hylobatidae |
| Nomascus leucogenys | Primates | Hylobatidae |
| Avahi betsileo | Primates | Indriidae |
| Avahi cleesei | Primates | Indriidae |
| Avahi laniger | Primates | Indriidae |
| Avahi meridionalis | Primates | Indriidae |
| Avahi occidentalis | Primates | Indriidae |
| Avahi peyrierasi | Primates | Indriidae |
| Avahi ramanantsoavanai | Primates | Indriidae |
| Avahi unicolor | Primates | Indriidae |
| Indri indri | Primates | Indriidae |
| Propithecus candidus | Primates | Indriidae |
| Propithecus coquereli | Primates | Indriidae |
| Propithecus coronatus | Primates | Indriidae |
| Propithecus deckenii | Primates | Indriidae |
| Propithecus diadema | Primates | Indriidae |
| Propithecus edwardsi | Primates | Indriidae |
| Propithecus perrieri | Primates | Indriidae |
| Propithecus tattersalli | Primates | Indriidae |
| Propithecus verreauxi | Primates | Indriidae |
| Eulemur albifrons | Primates | Lemuridae |
| Eulemur cinereiceps | Primates | Lemuridae |
| Eulemur fulvus | Primates | Lemuridae |
| Eulemur mongoz | Primates | Lemuridae |
| Eulemur rubriventer | Primates | Lemuridae |
| Eulemur rufifrons | Primates | Lemuridae |
| Eulemur rufus | Primates | Lemuridae |
| Hapalemur aureus | Primates | Lemuridae |
| Hapalemur griseus | Primates | Lemuridae |

| | | |
|---|---|---|
| Hapalemur meridionalis | Primates | Lemuridae |
| Hapalemur occidentalis | Primates | Lemuridae |
| Lemur catta | Primates | Lemuridae |
| Prolemur simus | Primates | Lemuridae |
| Varecia variegata | Primates | Lemuridae |
| Lepilemur ahmansonorum | Primates | Lepilemuridae |
| Lepilemur ankaranensis | Primates | Lepilemuridae |
| Lepilemur betsileo | Primates | Lepilemuridae |
| Lepilemur grewcockorum | Primates | Lepilemuridae |
| Lepilemur hollandorum | Primates | Lepilemuridae |
| Lepilemur hubbardorum | Primates | Lepilemuridae |
| Lepilemur jamesorum | Primates | Lepilemuridae |
| Lepilemur leucopus | Primates | Lepilemuridae |
| Lepilemur microdon | Primates | Lepilemuridae |
| Lepilemur milanoii | Primates | Lepilemuridae |
| Lepilemur mustelinus | Primates | Lepilemuridae |
| Lepilemur otto | Primates | Lepilemuridae |
| Lepilemur randrianasoloi | Primates | Lepilemuridae |
| Lepilemur seali | Primates | Lepilemuridae |
| Lepilemur wrightae | Primates | Lepilemuridae |
| Palaeopropithecus ingens | Primates | Palaeopropithecidae |
| Cacajao calvus | Primates | Pitheciidae |
| Xenothrix mcgregori | Primates | Pitheciidae |
| Elephas maximus | Proboscidea | Elephantidae |
| Loxodonta africana | Proboscidea | Elephantidae |
| Calomys hummelincki | Rodentia | Calomyscidae |
| Castor fiber | Rodentia | Castoridae |
| Chinchilla chinchilla | Rodentia | Chinchillidae |
| Chinchilla lanigera | Rodentia | Chinchillidae |
| Lagostomus crassus | Rodentia | Chinchillidae |
| Handleyomys rhabdops | Rodentia | Cricetidae |
| Juliomys rimofrons | Rodentia | Cricetidae |
| Juscelinomys candango | Rodentia | Cricetidae |
| Kunsia fronto | Rodentia | Cricetidae |
| Megalomys desmarestii | Rodentia | Cricetidae |
| Megalomys luciae | Rodentia | Cricetidae |
| Megaoryzomys curioi | Rodentia | Cricetidae |
| Microtus bavaricus | Rodentia | Cricetidae |
| Nesoryzomys darwini | Rodentia | Cricetidae |
| Nesoryzomys indefessus | Rodentia | Cricetidae |
| Oligoryzomys victus | Rodentia | Cricetidae |
| Oryzomys antillarum | Rodentia | Cricetidae |
| Oryzomys nelsoni | Rodentia | Cricetidae |
| Pennatomys nivalis | Rodentia | Cricetidae |
| Reithrodontomys microdon | Rodentia | Cricetidae |
| Rheomys mexicanus | Rodentia | Cricetidae |
| Thomasomys incanus | Rodentia | Cricetidae |
| Allactaga tetradactyla | Rodentia | Dipodidae |
| Boromys offella | Rodentia | Echimyidae |

| | | |
|---|---|---|
| Boromys torrei | Rodentia | Echimyidae |
| Brotomys voratus | Rodentia | Echimyidae |
| Capromys pilorides | Rodentia | Echimyidae |
| Geocapromys brownii | Rodentia | Echimyidae |
| Geocapromys columbianus | Rodentia | Echimyidae |
| Heteropsomys insulans | Rodentia | Echimyidae |
| Hexolobodon phenax | Rodentia | Echimyidae |
| Isolobodon montanus | Rodentia | Echimyidae |
| Isolobodon portoricensis | Rodentia | Echimyidae |
| Mesocapromys angelcabrerai | Rodentia | Echimyidae |
| Mesocapromys nanus | Rodentia | Echimyidae |
| Mysateles melanurus | Rodentia | Echimyidae |
| Phyllomys brasiliensis | Rodentia | Echimyidae |
| Plagiodontia aedium | Rodentia | Echimyidae |
| Plagiodontia ipnaeum | Rodentia | Echimyidae |
| Trinomys eliasi | Rodentia | Echimyidae |
| Myomimus roachi | Rodentia | Gliridae |
| Apodemus sylvaticus | Rodentia | Muridae |
| Apomys camiguinensis | Rodentia | Muridae |
| Apomys gracilirostris | Rodentia | Muridae |
| Conilurus albipes | Rodentia | Muridae |
| Conilurus capricornensis | Rodentia | Muridae |
| Coryphomys buehleri | Rodentia | Muridae |
| Crateromys heaneyi | Rodentia | Muridae |
| Hapalomys longicaudatus | Rodentia | Muridae |
| Leporillus apicalis | Rodentia | Muridae |
| Leporillus conditor | Rodentia | Muridae |
| Melomys aerosus | Rodentia | Muridae |
| Melomys fraterculus | Rodentia | Muridae |
| Mesembriomys gouldii | Rodentia | Muridae |
| Mesembriomys macrurus | Rodentia | Muridae |
| Mus fragilicauda | Rodentia | Muridae |
| Mus musculus | Rodentia | Muridae |
| Nesoromys ceramicus | Rodentia | Muridae |
| Niviventer cremoriventer | Rodentia | Muridae |
| Notomys alexis | Rodentia | Muridae |
| Notomys amplus | Rodentia | Muridae |
| Notomys cervinus | Rodentia | Muridae |
| Notomys fuscus | Rodentia | Muridae |
| Notomys longicaudatus | Rodentia | Muridae |
| Notomys macrotis | Rodentia | Muridae |
| Notomys mitchellii | Rodentia | Muridae |
| Notomys mordax | Rodentia | Muridae |
| Notomys robustus | Rodentia | Muridae |
| Otomys lacustris | Rodentia | Muridae |
| Palawanomys furvus | Rodentia | Muridae |
| Paulamys naso | Rodentia | Muridae |
| Praomys degraaffi | Rodentia | Muridae |
| Pseudomys albocinereus | Rodentia | Muridae |

| | | |
|---|---|---|
| Pseudomys australis | Rodentia | Muridae |
| Pseudomys bolami | Rodentia | Muridae |
| Pseudomys chapmani | Rodentia | Muridae |
| Pseudomys fieldi | Rodentia | Muridae |
| Pseudomys glaucus | Rodentia | Muridae |
| Pseudomys gouldii | Rodentia | Muridae |
| Pseudomys nanus | Rodentia | Muridae |
| Pseudomys novaehollandiae | Rodentia | Muridae |
| Pseudomys occidentalis | Rodentia | Muridae |
| Pseudomys oralis | Rodentia | Muridae |
| Pseudomys shortridgei | Rodentia | Muridae |
| Rattus hainaldi | Rodentia | Muridae |
| Rattus macleari | Rodentia | Muridae |
| Rattus nativitatis | Rodentia | Muridae |
| Rattus norvegicus | Rodentia | Muridae |
| Rattus praetor | Rodentia | Muridae |
| Rattus ranjiniae | Rodentia | Muridae |
| Rattus rattus | Rodentia | Muridae |
| Rattus satarae | Rodentia | Muridae |
| Rattus stoicus | Rodentia | Muridae |
| Rattus villosissimus | Rodentia | Muridae |
| Solomys sapientis | Rodentia | Muridae |
| Tokudaia muenninki | Rodentia | Muridae |
| Tokudaia osimensis | Rodentia | Muridae |
| Zyzomys pedunculatus | Rodentia | Muridae |
| Brachytarsomys albicauda | Rodentia | Nesomyidae |
| Brachytarsomys villosa | Rodentia | Nesomyidae |
| Brachyuromys betsileoensis | Rodentia | Nesomyidae |
| Brachyuromys ramirohitra | Rodentia | Nesomyidae |
| Eliurus carletoni | Rodentia | Nesomyidae |
| Eliurus grandidieri | Rodentia | Nesomyidae |
| Eliurus majori | Rodentia | Nesomyidae |
| Eliurus minor | Rodentia | Nesomyidae |
| Eliurus myoxinus | Rodentia | Nesomyidae |
| Eliurus petteri | Rodentia | Nesomyidae |
| Eliurus tanala | Rodentia | Nesomyidae |
| Eliurus webbi | Rodentia | Nesomyidae |
| Hypogeomys antimena | Rodentia | Nesomyidae |
| Macrotarsomys ingens | Rodentia | Nesomyidae |
| Monticolomys koopmani | Rodentia | Nesomyidae |
| Mystromys albicaudatus | Rodentia | Nesomyidae |
| Nesomys audeberti | Rodentia | Nesomyidae |
| Nesomys lambertoni | Rodentia | Nesomyidae |
| Nesomys rufus | Rodentia | Nesomyidae |
| Voalavo antsahabensis | Rodentia | Nesomyidae |
| Voalavo gymnocaudus | Rodentia | Nesomyidae |
| Marmota vancouverensis | Rodentia | Sciuridae |
| Petinomys genibarbis | Rodentia | Sciuridae |
| Petinomys setosus | Rodentia | Sciuridae |

| | | |
|---|---|---|
| Petinomys vordermanni | Rodentia | Sciuridae |
| Pteromyscus pulverulentus | Rodentia | Sciuridae |
| Sciurus vulgaris | Rodentia | Sciuridae |
| Spermophilus citellus | Rodentia | Sciuridae |
| Urocitellus brunneus | Rodentia | Sciuridae |
| Tupaia palawanensis | Scandentia | Tupaiidae |
