## Supplementary material for "Drivers of change in the realised climatic niche of terrestrial mammal species": Table S2; Table S3; Table S4; Table S5; Table S6; Figure S1; Figure S2

### SUPPLEMENTARY ONLINE MATERIALS

***Table S1*** *A list of species analysed (and their taxonomy), is provided as a separate spreadsheet.*

***Table S2*** *Performance of the multinomial model for predicting categories of climatic niche change, under alternative past climates: mid-Holocene under global circulation model IPSL-CM5A-LR (MID/IP); mid-Holocene under global circulation model MPI-ESM-P (MID/ME); last interglacial (LIG). For each model, three levels of tolerance were used to separate stability from change: 5%, 10%, 20%. The following stats are reported: Nagelkerke pseudo-$R^2$, overall percentage of correctly classified species (PCC), class-averaged percentage of correctly classified species, number (and proportion in parentheses) of species in each class of niche change.*

| | psd-$R^2$ | PCC (overall) | PCC (class) | N species shrink | N species shift | N species stable | N species expand |
|---|---|---|---|---|---|---|---|
| **MID/IP 5% tolerance** | 0.33 | 0.68 | 0.29 | 402 (68%) | 76 (13%) | 1 (0%) | 110 (19%) |
| **MID/IP 10% tolerance** | 0.38 | 0.63 | 0.33 | 355 (60%) | 147 (25%) | 15 (3%) | 72 (12%) |
| **MID/IP 20% tolerance** | 0.47 | 0.59 | 0.43 | 289 (49%) | 165 (28%) | 88 (15%) | 47 (8%) |
| **MID/ME 5% tolerance** | 0.25 | 0.73 | 0.24 | 438 (74%) | 62 (11%) | 4 (1%) | 85 (14%) |
| **MID/ME 10% tolerance** | 0.29 | 0.61 | 0.27 | 376 (64%) | 129 (22%) | 20 (34%) | 64 (11%) |
| **MID/ME 20% tolerance** | 0.38 | 0.51 | 0.36 | 287 (49%) | 167 (28%) | 104 (18%) | 31 (5%) |
| **LIG 5% tolerance** | 0.19 | 0.68 | 0.26 | 411 (70%) | 7 (12%) | 1 (0%) | 107 (18%) |
| **LIG 10% tolerance** | 0.28 | 0.62 | 0.36 | 371 (63%) | 137 (23%) | 11 (2%) | 70 (12%) |
| **LIG 20% tolerance** | 0.39 | 0.52 | 0.37 | 299 (51%) | 159 (27%) | 90 (15%) | 41 (7%) |

***Table S3*** *Validation of the multinomial model based on mid-Holocene climate (under global circulation model IPSL-CM5A-LR) and a tolerance of 20% to separate niche change from niche stability. Three validation metrics are represented: percentage of correctly classified species (PCC), where the observed niche class has the highest predicted probability; mean rank of the observed class, based on its predicted probability compared to other classes; mean difference in predicted probability of the most likely class vs the observed class. All metrics are calculated across all species, and separately for species within each class.*

| Metric | Overall | Shrink | Shift | Stable | Expand |
|---|---|---|---|---|---|
| PCC | 0.59 | 0.77 | 0.51 | 0.44 | 0 |
| rank | 1.64 | 1.27 | 1.66 | 1.98 | 3.28 |
| prediction difference | 0.15 | 0.06 | 0.17 | 0.18 | 0.52 |

***Table S4*** *Effect of utilising phylogenetic eigenvectors as predictors of niche change in the multinomial model, based on mid-Holocene climate (under global circulation model IPSL-CM5A-LR) and a tolerance of 20% to separate niche change from niche stability. For models using an incremental number of eigenvectors, the pseudo-$R^2$ (Nagelkerke), degrees of freedom, and AIC values are reported.*

| **n. eigenvectors** | **Pseudo-$R^2$** | **df** | **AIC** |
|---|---|---|---|
| 0 | 0.47 | 57 | 1192 |
| 5 | 0.49 | 72 | 1204 |
| 10 | 0.53 | 87 | 1192 |
| 20 | 0.57 | 117 | 1204 |

***Table S5*** *Effect of phylogenetic uncertainty on model's classification, based on mid-Holocene climate (under global circulation model IPSL-CM5A-LR), and a tolerance of 20% to separate niche change from niche stability. The table reports the number of species predicted to be in the same niche class (most probable class) when using intrinsic traits imputed using 10 different phylogenies.*

| **N predictions with the same classification** | **N species** |
|---|---|
| 6/10 | 3 |
| 7/10 | 1 |
| 8/10 | 4 |
| 9/10 | 5 |
| 10/10 | 576 |

***Table S6*** *Statistical significance (p value) of the multinomial model's coefficients, for the model based on* the IPSL-CM5A-LR (IP) and the MPI-ESM-P (ME) general circulation models. *Symbols report p value levels, as: "+" p<0.1; "*" p<0.05; "**" p<0.01; "***" p<0.001.*

| **Variable** | **shift/shrink** | **stable/shrink** | **expand/shrink** | **shift/shrink** | **stable/shrink** | **expand/shrink** |
|---|---|---|---|---|---|---|
| **Climate (GCM)** | **IP** | **IP** | **IP** | **ME** | **ME** | **ME** |
| Intercept | 0.0000*** | 0.0000*** | 0.0000*** | 0.0032** | 0.0000*** | 0.0000*** |
| rlm_AT | 0.0000*** | 0.7414 | 0.0002*** | 0.1202 | 0.8617 | 0.0049** |
| rlm_ATPA | 0.9766 | 0.2820 | 0.2107 | 0.6186 | 0.5373 | 0.9273 |
| rlm_IMnOC | 0.0044** | 0.0001*** | 0.0002*** | 0.0395* | 0.0406* | 0.0001*** |
| rlm_IMPA | 0.1546 | 0.4779 | 0.0420* | 0.6423 | 0.0848+ | 0.8902 |
| rlm_multi_rlm | 0.8972 | 0.7032 | 0.0034** | 0.4420 | 0.0518+ | 0.0278* |
| rlm_NAnTRANSB | 0.5968 | 0.7898 | 0.3172 | 0.6048 | 0.9764 | 0.9031 |
| rlm_NT | 0.8398 | 0.5961 | NA | 0.0121* | 0.7385 | 0.1943 |
| rlm_PA | 0.7677 | 0.1478 | 0.8368 | 0.8238 | 0.2474 | 0.9130 |
| Clim_maint | 0.0000*** | 0.0000*** | 0.0225* | 0.1848 | 0.0000*** | 0.0300* |
| Diet.Plant | 0.2092 | 0.1101 | 0.8574 | 0.0374* | 0.0462* | 0.6935 |
| Diet.Vertebrate | 0.6635 | 0.8046 | 0.0676+ | 0.2515 | 0.5167 | 0.3550 |
| Diet.Invertebrate | 0.0994+ | 0.0985+ | 0.0647+ | 0.0012** | 0.0174* | 0.1362 |
| Mass.g | 0.0607+ | 0.1459 | 0.4849 | 0.2583 | 0.2880 | 0.0672+ |
| GestLen_d | 0.2190 | 0.9394 | 0.0979+ | 0.5785 | 0.3712 | 0.0546+ |
| IntInt_d | 0.0181* | 0.9695 | 0.7043 | 0.0845+ | 0.7146 | 0.0347* |
| Human_popd_4kBC | 0.8200 | 0.0722+ | 0.9736 | 0.7355 | 0.0078** | 0.9844 |
| Land_use_4kBC | 0.8156 | 0.6569 | 0.6967 | 0.4970 | 0.9292 | 0.5659 |
| Human_popd_2017 | 0.4452 | 0.0206* | 0.8554 | 0.5448 | 0.1340 | 0.7674 |
| Land_use_2017 | 0.0376* | 0.9935 | 0.3190 | 0.1599 | 0.7511 | 0.0499* |

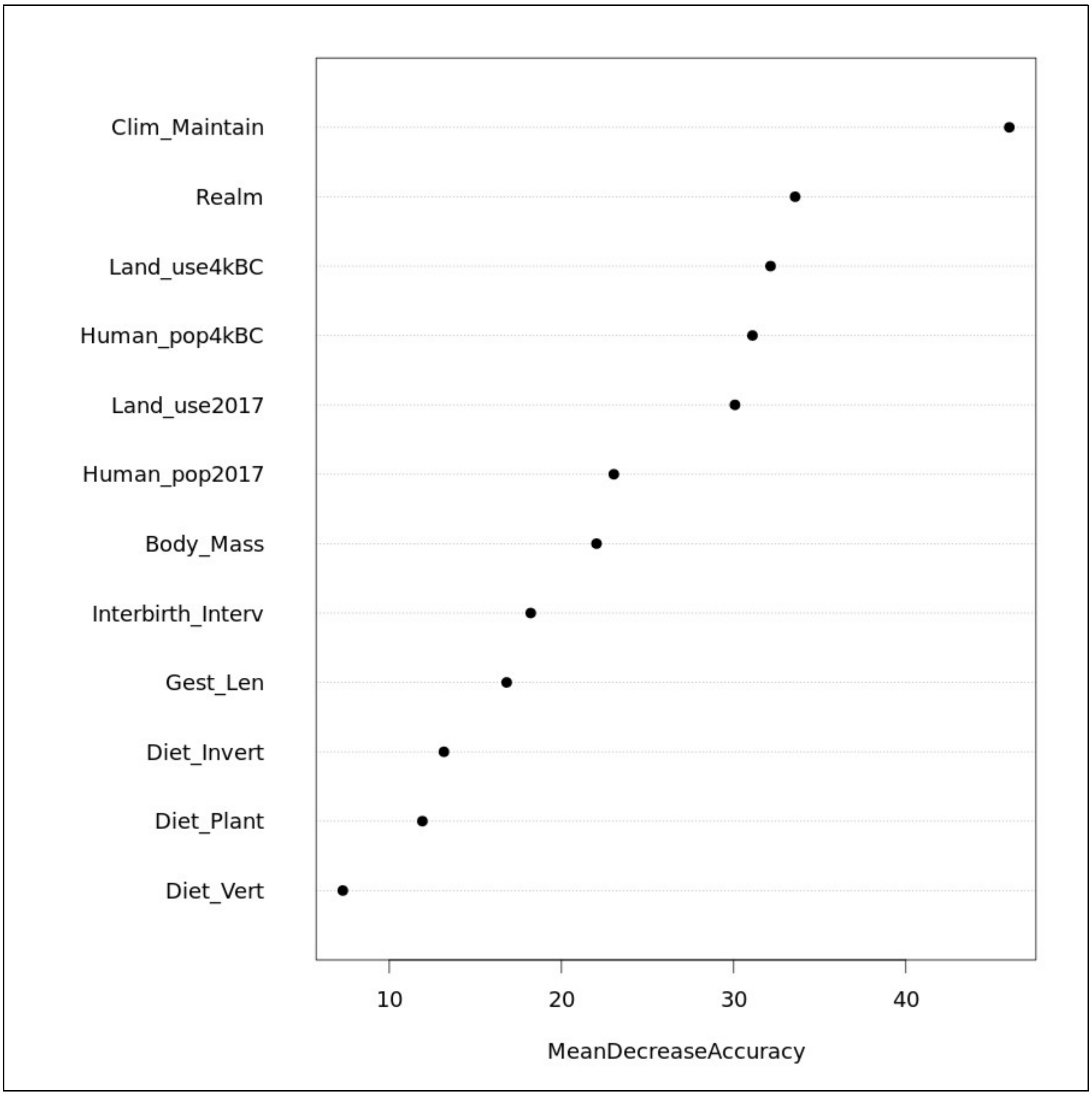


***Figure S1*** *Importance of predictor variables in a random forest model to classify species' niche classes. The measure of importance (x-axis) represents variables' ability to influence model's classification.*

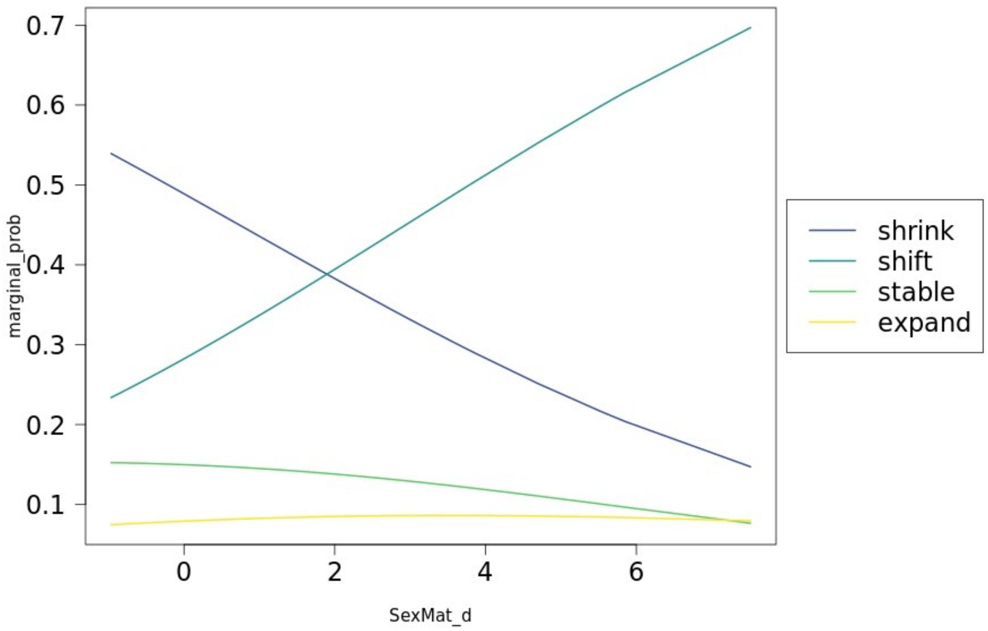


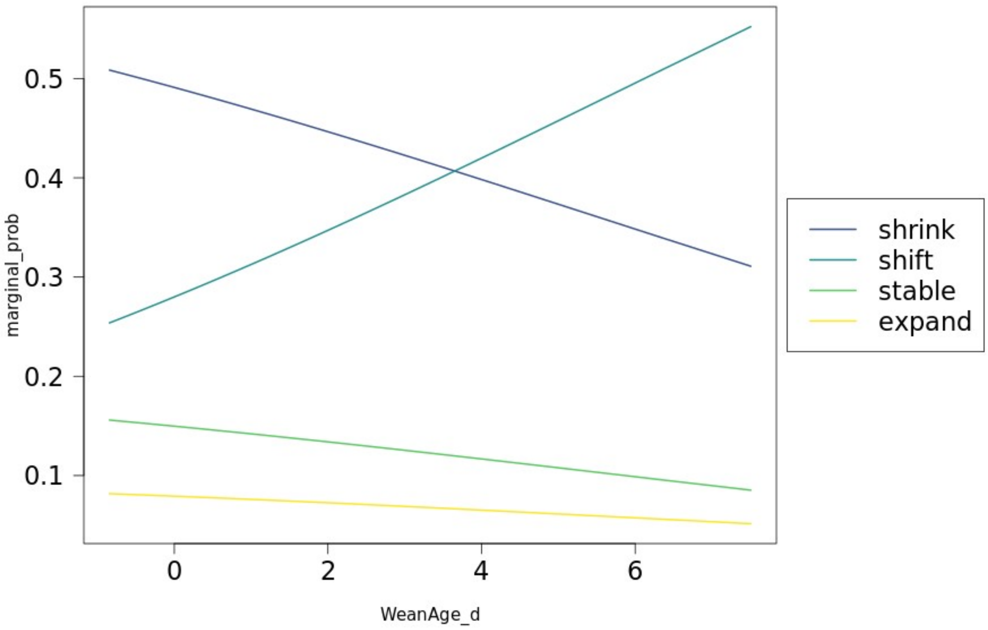


***Figure S2*** *Partial effect plot of the relationships between age at sexual maturity and weaning age (scaled for the multinomial model) and the probability of species to be assigned to one of four categories of niche change (shrink, stable, shift, expand).*
